# Development of a Self-Report Measure of Reward Sensitivity: A Test in Current and Former Smokers

**DOI:** 10.1101/128793

**Authors:** John R Hughes, Peter W Callas, Jeff S Priest, Jean-Francois Etter, Alan J Budney, Stacey C Sigmon

## Abstract

**Introduction:** Tobacco use or abstinence may increase or decrease reward sensitivity. Most existing measures of reward sensitivity were developed decades ago, and few have undergone extensive psychometric testing.

**Methods:** We developed a 58-item survey of the anticipated enjoyment from, wanting for, and frequency of common rewards (the Rewarding Events Inventory – REI). The current analyses focuses on ratings of anticipated enjoyment. The first validation study recruited current and former smokers from internet sites. The second study recruited smokers who wished to quit and monetarily reinforced them to stay abstinent in a laboratory study, and a comparison group of former smokers. In both studies, participants completed the inventory on two occasions, 3-7 days apart. They also completed four anhedonia scales and a behavioral test of reduced reward sensitivity.

**Results:** Half of the enjoyment ratings loaded on four factors: socializing, active hobbies, passive hobbies, and sex/drug use. Cronbach alpha coefficients were all ≥ 0.73 for overall mean and factor scores. Test-retest correlations were all ≥ 0.83. Correlations of the overall and factor scores with frequency of rewards, anhedonia scales were 0.19 – 0.53, except for the sex/drugs factor. The scores did not correlate with behavioral tests of reward and did not differ between current and former smokers. Lower overall mean enjoyment score predicted a shorter time to relapse.

**Discussion:** Internal reliability and test-retest reliability of the enjoyment outcomes of the REI are excellent, and construct and predictive validity are modest but promising. The REI is comprehensive and up-to-date, yet is short enough to use on repeated occasions. Replication tests, especially predictive validity tests, are needed.

**Implications:** Both use of and abstinence from nicotine appears to increase or decrease how rewarding non-drug rewards are; however, self-report scales to test this have limitations. Our inventory of enjoyment from 58 rewards appears to be reliable and valid as well as comprehensive and up-to-date, yet is short enough to use on repeated occasions. Replication tests, especially of the predictive validity of our scale, are needed.

## 1. INTRODUCTION

Several lines of evidence suggest that nicotine use or abstinence can increase, decrease, or not change the efficacy of non-drug rewards ^1, 2^. In addition, a central theme in many treatments for drug abuse is an attempt to increase sensitivity to non-drug rewards ^3, 4^. Reward sensitivity can be measured by behavioral tests, neuroimaging tests, and self-report scales. Behavioral and neuroimaging tests most often focus on operant measures of reward seeking, whereas self-report measures mostly focus on enjoyment from rewards ^5^. There are many (>21) such self-report measures ^6,5,7,8^. These scales typically ask how pleasurable several rewards would be for an individual. The existing scales are often long (survey > 150 rewards) ^9-11^, fail to ask about more recent rewards (e.g., some scales are > 40 years old) ^9, 10^, or have undergone limited psychometric testing. For example, one widely used scale is the Pleasant Events Scale (PES). This test has good psychometrics ^10^ but it is lengthy (640 questions, 45-60 minutes to complete) and since it was developed 40 years ago, does not ask about more recent rewards such as texting, social media, or internet browsing. The current paper describes a new self-report measure (The Rewarding Events Inventory-REI) that uses more current rewards, is comprehensive, but brief enough (58 questions) that it could be used on a repeated basis, and asks about more up-to-date possible rewards.

## 2. METHODS

### 2.1 Scale development

The REI was developed for use in a study on whether smoking cessation decreases reward sensitivity ^12^. We began by examining the 21 existing reward inventories, anhedonia scales, and apathy scales to obtain a list of commonly cited rewards. Next, we added newer rewards (e.g., browsing the internet) not included in these scales. This resulted in a list of 476 rewards. We then deleted rewards that we believed would occur rarely and categorized the rewards into specific themes (e.g., alcohol/other drug use, consumerism/shopping, and eating) to identify overlapping rewards. All decisions regarding inclusion of rewards were made via consensus of the authors. One challenge was whether questions should refer to a) past rewards, b) current rewards, c) “usual” rewards, or d) future (anticipated or hypothetical) rewards ^13, 14^. We chose to ask about anticipated rewards because they are probably of greater clinical significance than past rewards ^15,16^, plus it allows ratings of rewards that are infrequent or have never occurred. We decided to use broad rather than specific descriptions (“sports” vs skiing, basketball, etc), to obtain adequate incidence rates.

This process resulted in 155 rewards. The authors then rated the 155 rewards on enjoyment, wanting, and frequency, as well as clarity. Based on the magnitude, clarity, overlap, and floor/ceiling effects from these ratings, we reduced the number of rewards to 99. Next, to better sample young adults we asked 20 young adults (18-24 years old) to record on a website at least five rewards that happened in the previous week on two consecutive weeks. This resulted in no additions, but, did result in two revisions to the existing list of rewards.

We initially developed three response options about the 99 rewards: i.e., how much participants enjoyed each reward, how much they wanted it, and how often it occurred. We asked about wanting vs enjoyment because animal research suggests these are different behavioral states ^17, 18^. However, although indirect measures can dissociate wanting from enjoying in humans, when asked to rate both wanting and enjoyment humans rarely distinguish between the two ^17, 18^. Consistent with this, we found a very high correlation between enjoyment and wanting, and very few instances of discordances between the two. Also, participants in our pilot work appeared to have more difficulty rating wanting than enjoyment. We also noted that there were often discrepancies between the enjoyment and frequency ratings because many factors other than enjoyment; e.g. availability, influence the frequency of rewards. For the above reasons, the current analyses were based solely on the enjoyment ratings. To assess enjoyment, the REI asked participants to “rate how much you would enjoy each reward using the following categories: “I would extremely enjoy it, I would enjoy it a lot, I would enjoy it some, I would enjoy it a little, I would not enjoy it”.

### 2.2 Validation Studies

We used results from two studies to examine the psychometrics of the REI. Both the development work and these two studies were approved by the University of Vermont Committees on the Use of Human Subjects in Research.

In the first study, we sent invitations via emails to current or former smokers who had visited a stop smoking website (www.stop-tabac.ch) developed by one of the authors (JFE). These participants had previously volunteered to participate in surveys without monetary reimbursement. We also posted links on other websites such as stopsmokingcenter.net and virtualmedicalcentre.com. Inclusion criteria were a) English is native language, b) > 18 yrs old, c) current or past daily smoker, and d) no current psychiatric or neurological problem that could influence reward processes (e.g. Parkinson’s or depression). The website had participants complete the survey on three occasions over approximately one week.

The second study was an experimental test of whether smoking cessation decreases reward sensitivity that is described in a separate paper in this special issue of NTR^12^. During the first week, current smokers smoked as usual, and during the last 4 weeks they were reimbursed to remain abstinent. Smokers completed the REI scale and several other measures twice/week. For the current analysis we used only the data from the two visits in the first week when smokers were still smoking. The study also included former smokers who completed the REI four times over 2 weeks; again, we used their first two surveys.

We collected several outcomes to test construct validity of the enjoyment ratings: a) frequency of rewards subscale of the REI, by asking participants to “rate how often the reward has occurred in the last week” from “It occurred every day in the last week, on most days in the last week, on a few days in the last week, on one day in the last week, did not occur in the last week.”, b) a behavioral measure of decreased reward sensitivity - the Effort Expenditure for Rewards Task (EEfRT) - that examines responding as a function of response cost, reward magnitude and probability of reward^19^, c) two anhedonia scales: the Apathy Evaluation Scale (AES) and the Temporal Experience of Pleasure Scale (TEPS)^14,5,7,8^, and d) a measure of positive affect (PA) via the Positive and Negative Affect Scale (PANAS) ^20^. The major inclusion criteria were the same as the first study except this study required smoking > 10 cigarettes/day currently or in the past, and current smokers had to be trying to quit.

We pooled the results of the two studies for two reasons. First, factor analysis requires large sample sizes, especially when testing >50 items^21^. Second, combining studies increased the range of demographics and smoking history outcomes. Exploratory analyses suggests the results were very similar for current vs former smokers and for Study 1 vs Study 2. The 440 participants were middle aged, and mostly White/non-Latinos with some college education. About half were women and, among current smokers, half smoked more than 20 cigarettes/day (Table 1).

**Table 1.** Demographics and Smoking History of Participants.

|  | Study 1 | Study 2 | Current<br>Smokers | Former<br>Smokers |
| --- | --- | --- | --- | --- |
|  | (*n* = 162) | (*n* = 278) | (*n* = 269) | (*n* = 171) |
| Age (*M* ± *SD*) | 48 ± 12 | 42 ± 15*** | 41 ± 14 | 49 ± 13*** |
| % Women | 60% | 45%** | 48% | 54% |
| % White/Non-Latino | 90% | 86% | 87% | 89% |
| % Some college or more | 81% | 73%* | 69% | 86%*** |
| Cigarettes per day |  |  |  |  |
| 1-9 cigarettes/day | 31% | 0% | 7% | — |
| 10-19 cigarettes/day | 29% | 49% | 44% | — |
| 20 cigarettes/day | 19% | 30% | 28% | — |
| > 20 cigarettes/day | 21% | 20% | 20% | — |

M = mean, SD = standard deviation. Categorical variables were tested using the Pearson chi-square; Continuous variables using the Wilcoxon Rank Sum Test \**p*<.05; \*\**p*<.01; \*\*\**p*<.001

### 2.3. Data Analysis

After initial inspection of the data from Study 1, we deleted 41 rewards due to a high incidence, of “don’t know/unclear responses,” very low or very high enjoyment rating (to avoid floor and ceiling effects), high correlation with another reward, or very low frequency of occurrence. When different orders of questions were used, there was no difference in results for the 10 rewards at the beginning or end of the scales, suggesting significant response fatigue did not occur. For the remaining 58 rewards, we examined a) factor structure, b) internal reliability via Cronbach’s alpha, c) test-retest validity by comparing scores between the first two sessions of each study, d) construct validity by comparing ratings of enjoyment with ratings of the frequency of rewards and to the EEfRT, AES, TEPS and PANAS PA scores, and d) predictive construct validity by testing whether the REI differed between current and former smokers, and whether baseline REI scores predicted time to relapse among current smokers trying to quit. We conducted several statistical tests and, thus, some of our results may be false positives. We did not correct for p values because many statisticians believe this is not appropriate in early research in an area ^22, 23^.

For the factor analysis, a polychoric correlation matrix was generated and used in the Factor 9.2 Program ^24^ to determine the number of factors to extract, based on parallel analysis and minimum rank factor analysis ^25^ Maximum likelihood estimates were then generated in SAS 9.4 (PROC FACTOR) (SAS Institute Inc., Cary, NC) using oblique promax rotation. We used relatively stringent criteria for determining factors. Rewards were placed with factors for which rotated loadings were ≥ 0.30. Rewards with loading <0.30 on all factors, loading ≥ 0.30 on more than one factor, or loading ≥ 0.30 on different factors for Visit 1 and Visit 2 were not included in any factor but were included in the overall mean reward score.

For each psychometric test, we examined outcomes both for the overall mean score and the factor scores of the enjoyment ratings. For internal reliability, we calculated Cronbach’s alpha. For test-retest reliability, we calculated Intraclass Correlation Coefficients. For construct validity we examined Pearson Product correlations between REI scores and EEfRT, reward frequency, AES, TEPS and PANAS scores. For predictive validity, we tested a) whether the REI scores differed between current and former smokers via a linear regression that included baseline differences in the groups as covariates, and b) whether, in the second study, the REI scores from the first week predicted the probability of relapse when smokers were trying to quit using a proportional hazards regression.

## 3. RESULTS

### 3.1 Introductory Remarks

The actual values for the REI, EEfRT, PANAS, TPS, and AES during the first week of the second study are reported in detail in the accompanying paper in this issue ^12^. Across the two visits, the mean enjoyment score (standard deviation) of the 58 rewards on a scale of 1 = I would not enjoy it” to 5 “I would extremely enjoy” was 3.6 (0.5) for both visits. The three highest rated rewards were “go on vacation” (4.5), “be told I am loved” (4.4), and “kiss someone romantically” (4.3). The three lowest scores were “use marijuana or other drugs” (1.6), “watch sports” (2.5) and “drink alcohol” (2.6). When we posted the 58 reward REI Scale on a website (www.stop-tabac.ch), a new sample of 157 respondents took a median of 4.3 minutes (Interquartile range = 3.4-6.0 minutes) to complete the enjoyment scale.

### 3.2 Factor Analysis

Half of the enjoyment ratings (29) loaded onto four factors that we labeled “socializing”, “active hobbies”, “passive hobbies”, and “sex/drug use” (Appendix Table 1). The loadings for these rewards were very similar for Visits 1 and 2. Several other rewards loaded on a fifth factor but item loading on this factor was not consistent between Visit 1 and Visit 2. The four factors included were moderately inter-correlated (r = ·26-.55 for Visit 1 and ·24-.55 for Visit 2). The mean enjoyment scores for the socializing, active hobbies, and passive hobby factor scores ranged from 3.5-3.6 (sd = 0.5-0.8) across the factors and visits. The mean scores for the sex/drug use scores for both visits were 3.1 (0.8).

### 3.3 Reliability

Reliability analyses were based on the first two sessions in both studies. Cronbach’s alphas were all > 0.70; i.e. indicating “moderate” to “excellent” reliability ^26^ (Table 2). Intraclass coefficients of test-retest stability across the overall mean and the three factors were all > 0.83; i.e. “excellent” (Table 2).

**Table 2.** Internal Reliability (n=440), Test-Retest Reliability (n = 348) and Concurrent Validity (n=278)^a^

|  |  | Overall Mean<br>Score | Socializing | Active<br>Hobbies | Passive<br>Hobbies | Sex/Drug<br>Use |
| --- | --- | --- | --- | --- | --- | --- |
| **Internal Reliability** |  |  |  |  |  |  |
| Cronbach's alpha |  |  |  |  |  |  |
| Visit 1 |  | 0.94 | 0.88 | 0.83 | 0.72 | 0.71 |
| Visit 2 |  | 0.95 | 0.90 | 0.84 | 0.70 | 0.73 |
| **Test-Retest Reliability** |  |  |  |  |  |  |
| ICC<sup>b</sup> |  | 0.89 | 0.89 | 0.87 | 0.83 | 0.88 |
| **Concurrent Validity<sup>b</sup>** |  |  |  |  |  |  |
| Visit 1 | Frequency<sup>c</sup> | 0.52 | 0.44 | 0.44 | 0.40 | 0.24 |
|  | EEfRT | -0.11 | -0.14 | -0.06 | -0.01 | -0.01 |
|  | AES | -0.51 | -0.53 | -0.46 | -0.30 | -0.15 |
|  | TEPS | -0.53 | -0.44 | -0.45 | -0.40 | -0.25 |
|  | PANAS PA | 0.53 | 0.52 | 0.44 | 0.35 | 0.22 |
| Visit 2 | EEfRT | -0.03 | -0.07 | 0.03 | 0.06 | -0.02 |
|  | AES | -0.38 | -0.39 | -0.37 | -0.19 | -0.15 |
|  | TEPS | -0.49 | -0.44 | -0.41 | -0.35 | -0.19 |
|  | PANAS PA | 0.53 | 0.51 | 0.45 | 0.33 | 0.22 |

<sup>a</sup>AES = Apathy Evaluation Scale, EEfRT= Effort Expenditure for Rewards Task, ICC = Intraclass Correlation Coefficient, PANAS PA = Positive and Negative Affect Scale, Positive Affect Score, REI = Rewarding Events Inventory, TEPS = Temporal Experience of Pleasure Scale
p<0.01 for all correlations except: p<0.05 for AES and Sex/Drug Use and p > 0.05 for all EEfRT correlations
<sup>b</sup>Visit 1 vs Visit 2
<sup>c</sup>Administered only at Visit 1

### 3.4 Construct Validity

As expected, higher overall REI enjoyment score, as well as the socialization score and the active hobbies subscores, were correlated with a greater frequency of rewards, higher PANAS PA score, and lower AES and TEPS anhedonia scores (Table 2) (r = ·37-.53). The same was true for the passive hobby scores and sex/drugs scores but to a lesser degree (r= ·15-.40). The REI was not correlated with EEfRT scores.

### 3.5 Predictive Validity

Contrary to our prediction, overall enjoyment score and factor scores did not differ between current and former smokers (Table 3). Higher overall and factor scores did prospectively predict a lower probability of relapsing during the laboratory study (Table 4). For example, each one unit increase in the overall enjoyment score at Visit 1 decreased the probability of relapse by 27%.

**Table 3.** Adjusted Overall Mean REI and Factor Scores for Current (n = 269) vs. Former Smokers (n = 171)^a^

|  | Overall Mean |  | Socializing |  | Active Hobbies |  |
| --- | --- | --- | --- | --- | --- | --- |
|  | Current Smoker | Former Smoker | Current Smoker | Former Smoker | Current Smoker | Former Smoker |
| Visit 1 | 3.7 | 3.6 | 3.8 | 3.7 | 3.7 | 3.6 |
| Visit 2 | 3.6 | 3.6 | 3.7 | 3.7 | 3.6 | 3.6 |

|  | Passive Hobbies |  | Sex/Drug Use |  |
| --- | --- | --- | --- | --- |
|  | Current Smoker | Former Smoker | Current Smoker | Former Smoker |
| Visit 1 | 3.6 | 3.6 | 3.1 | 3.1 |
| Visit 2 | 3.5 | 3.7* | 3.1 | 3.1 |

\* p=.02, for current vs former smoker
<sup>a</sup>Adjusted for sex, race, and education

**Table 4.** Hazard Ratios for Time to Relapse

|  | Overall Score | Socializing | Active Hobbies | Passive Hobbies | Sex/drugs |
| --- | --- | --- | --- | --- | --- |
| Visit 1 | 0.73* | 0.94 | 0.84 | 0.83 | 0.85 |
| Visit 2 | 0.72** | 0.89 | 0.82* | 0.81* | 0.81** |

\* $p \leq .10$
\*\* $p \leq .05$

### 3.6. Moderators

Women scored higher than men on the overall enjoyment score and the socializing and passive hobby factor scores, but scored lower on the sex/drug use scores (Appendix Table). Older participants scored lower than younger participants on the overall mean score and all factors except for passive hobbies (which showed a similar trend). Ethnicity/race and education did not moderate scores

## 4. DISCUSSION

Our Rewarding Events Inventory (see Appendix for the final version) includes three outcomes: enjoyment from rewards, wanting of rewards and frequency of rewards. The psychometric analyses in this report focuses on the enjoyment ratings for the reasons cited above. The list of rewards in the REI appears to be comprehensive (includes 58 rewards) and up-to-date, yet the enjoyment scale of the REI can be completed by most participants within 5 minutes. Internal reliability and test-retest reliability of enjoyment ratings were excellent, concurrent validity was good, but predictive validity was unclear.

Our scale is most similar to the PES ^10^, the Pleasant Activities List ^11^, and the Reinforcement Survey Scale ^9^. Factor analyses of these scales suggested socializing, solitary, craft, and sexual factors which is similar to our analysis. Only the PES has had psychometric testing and our results are comparable to their results ^10^. Our scale may be preferable to these three scales for several reasons. First, these three scales have 2-4 times the number of rewards as our scale and take about 30-60 minutes to complete. Second, two of the scales were published in 1981-1982, and thus fail to include more recent rewards. Third, these scales ask about past enjoyment, whereas our scale asks about anticipated enjoyment. We focused on anticipated rewards because future behavior and much psychopathology are based on perceived outcomes.

Our study had limitations. First, the REI was not based on any specific theoretical conceptualization of anhedonia. Also, the REI measures only anticipatory anhedonia and not consummatory anhedonia; thus, the scale does not measure actual enjoyment when the reward occurs. This is important because anticipating and consuming rewards appear to be two different phenomena ^27^. Our use of convenience samples decreased our external validity, and our use of only current and former smokers may mean that our results may not generalize to never smokers. In addition, our sample had few minorities and few participants with a high school-only education. To conduct factor analyses, we had to combine results from two different studies, which, although increasing the range of possible scores, may have added unwanted variance.

We hope that publishing our scale will prompt researchers to conduct rigorous tests of the REI. Future studies especially need to include more stringent validity tests; e.g. whether scores differ in those with depression, schizophrenia, or drug withdrawal. Another important test would be whether the REI predicts outcomes, or whether it changes with clinical improvement. For example, the REI should change with successful implementation of contingency management ^3^ or behavioral activation therapies ^28^, or with certain medications; e.g., antidepressants ^29^. In addition, our decision to focus only on anticipated enjoyment was based on our anecdotal experience and clinical logic. Delineation of the relationships among enjoyment of, wanting for, and frequency of rewards is clearly indicated. Our REI scale includes questions about wanting and frequency as well as enjoyment so that future researchers can examine these relationships.

In summary, we have developed what we believe is a comprehensive, up-to-date, yet brief inventory, that can be used to measure self-reported reward sensitivity on a repeated basis. In addition, it is one of the few scales that has been shown to have test-retest and prospective validity. Replication of our results in more generalizable samples and tests of the clinical utility of our scale are necessary prior to its widespread use, and we encourage such tests.

## Contributors

JRH, AJB and PWC designed the study and obtained funding. JRH, AJB, JRF, JFE, PWC and SCS conducted the study. PC, JP and JH supervised data analysis. All authors participated in data analysis as well as interpretation and writing of the manuscript.

## Acknowledgements

We thank Michael Treadway for sharing his EEfRT task with us. We also thank Bonita Basnyat, Erin Kretzer, Doris Gasangwa, Grey Norton, Stanley See, and Hao Yani for assistance in conducting the study and Jessie McNabb for help in preparing the manuscript.

## APPENDIX Rewarding Events Inventory

### Instructions

The REI asks participants to rate 58 common rewards on three outcomes: enjoying, wanting or frequency. We found enjoying and wanting to be highly correlated and that participants stated rating enjoying was much easier than rating wanting. In addition, we found many discrepancies between enjoyment and frequency of reward, probably because other factors (e.g., availability of the reward) influence the frequency of rewards. In summary, we believe the enjoyment ratings are better measures of reward sensitivity than the wanting of frequency ratings; thus, if, due to response burden or time concerns, researchers can only use one scale, we suggest it be the enjoyment scale. On the other hand, we encourage researchers to ask all three outcomes to help understand the relationship among enjoyment of, wanting for, and frequency of rewards.

The participant instructions for the three outcomes is listed below. We suggest not asking participants to rate enjoyment, wanting and frequency for a reward at the same time because this may cause a false concordance among these three response options. Instead, we suggest participants first rate all rewards on one of the three outcomes and then move on to rating all rewards on another outcome. One can randomize participants to order of outcomes being assessed and to order of questions.

### Participant Instructions

In the sections that follow you will be asked to review a list of rewards three times. First on how much you would want it. Second, how much you would enjoy it. Third, how frequently it has occurred in the last week. At the beginning of each section, you will be given more detailed instructions.

#### Enjoying

Rate how much you would enjoy each reward. Please note that “Enjoying” is not the same as “Wanting.” It is possible to enjoy something even though you don’t want it enough to put any time, effort or money into experiencing it. In this section, please tell us how much you would ENJOY the item.

Response choices:

- I would extremely enjoy it
- I would enjoy it a lot
- I would enjoy it some
- I would enjoy it a little
- I would NOT enjoy it

#### Wanting

On the following questions rate how much you would want each reward to occur. Please note that “Wanting” is different than “Enjoying.” In this section we are interested in wanting-that is, how much would you be willing to spend time, money, or effort to be able to experience it?

Response choices:

- I would extremely want it
- I would want it a lot
- I would want it some
- I would want it a little
- I would NOT want it

#### Frequency

Rate how often the reward has occurred to you in the last week. Response choices:

- It occurred every day in the last week
- It occurred on most days in the last week
- It occurred on a few days in the last week
- It occurred on one day in the last week
- It did NOT occur in the last week

#### Rewards

1. Give a party or get-together
2. Meet someone new
3. Talk on the telephone
4. Do art-related work
5. Give gifts; do favors for people
6. Reminisce, talk about old times
7. Solve a puzzle, crossword, etc
8. Text, email, or chat on the internet
9. Celebrate holidays / birthdays
10. Be told that I am loved
11. Take a bike ride
12. Be alone
13. Watch sports
14. Surf the internet
15. Smoke tobacco
16. Express my love to someone
17. Do craft work: sewing, woodworking, etc
18. Drive a car, motorcycle, etc
19. Hear a good joke
20. Sit and think; have daydreams
21. Watch movies
22. Have a meal or snack with friends
23. Engage in sexual activity
24. Go to a party or other social event
25. Take a stay at home vacation
26. Receive a compliment
27. Play games (board, card, computer, video, etc)
28. Play a sport
29. Do gardening or yard work
30. Go on a vacation
31. Attend a performance: concert, play, etc
32. Work on home improvements
33. Help someone
34. Make a new friend
35. Take a nap
36. Listen to music
37. Talk about sex
38. Watch people
39. Have spare time
40. Get mail or email from friends or family
41. Plan trips or vacations
42. Eat snacks
43. Start a new project
44. Eat a meal out
45. Watch TV
46. Go to a bar, tavern, club, etc
47. Go shopping
48. Drink alcohol
49. Use marijuana or other drugs
50. Do activities with a friend
51. Kiss someone romantically
52. Cook
53. Take a walk
54. Do great in my classes or at work
55. Read for pleasure
56. Be with a pet or other animals
57. Be popular at a gathering
58. Be outdoors in nature

**Appendix Table 1.**
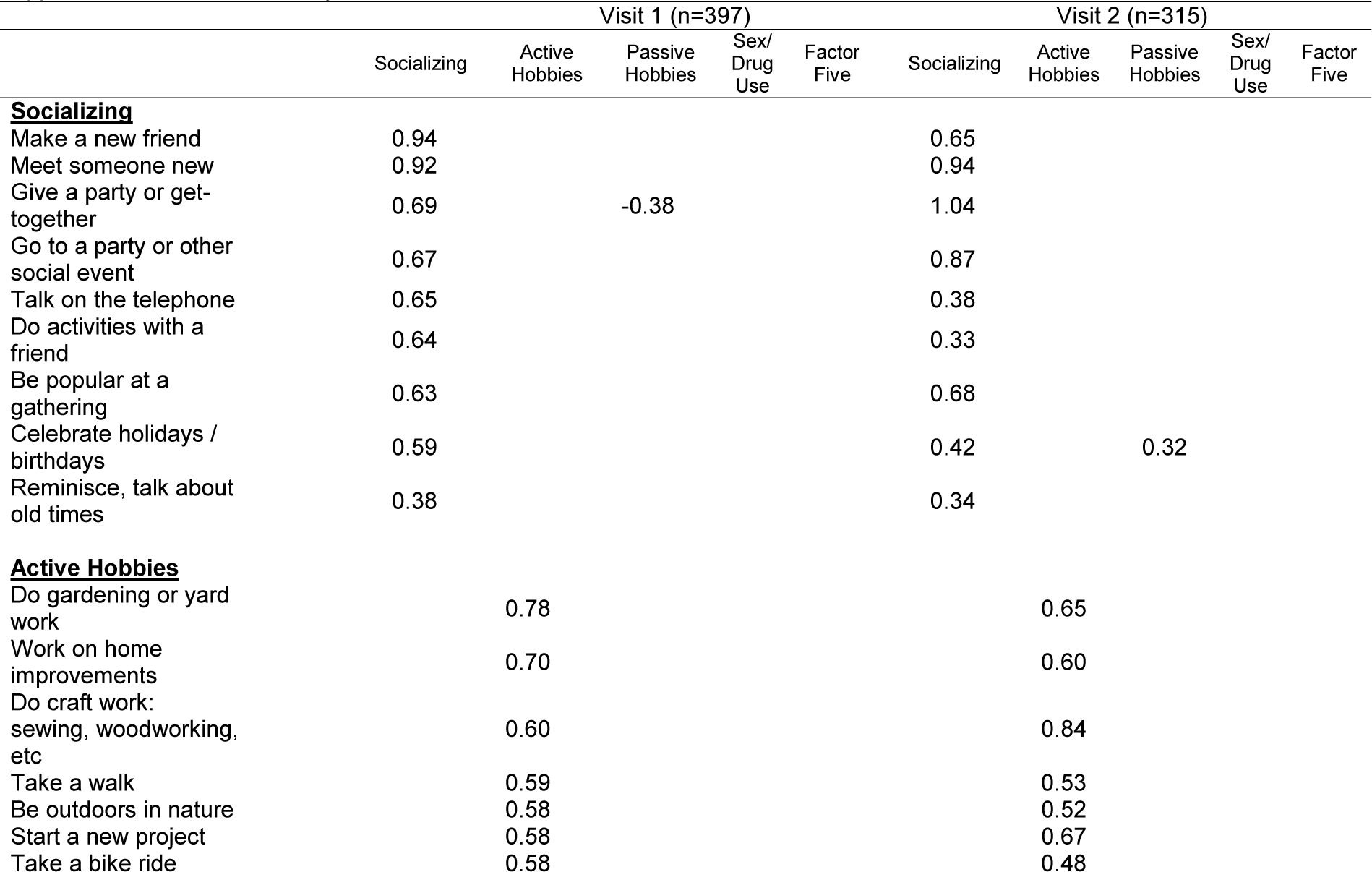
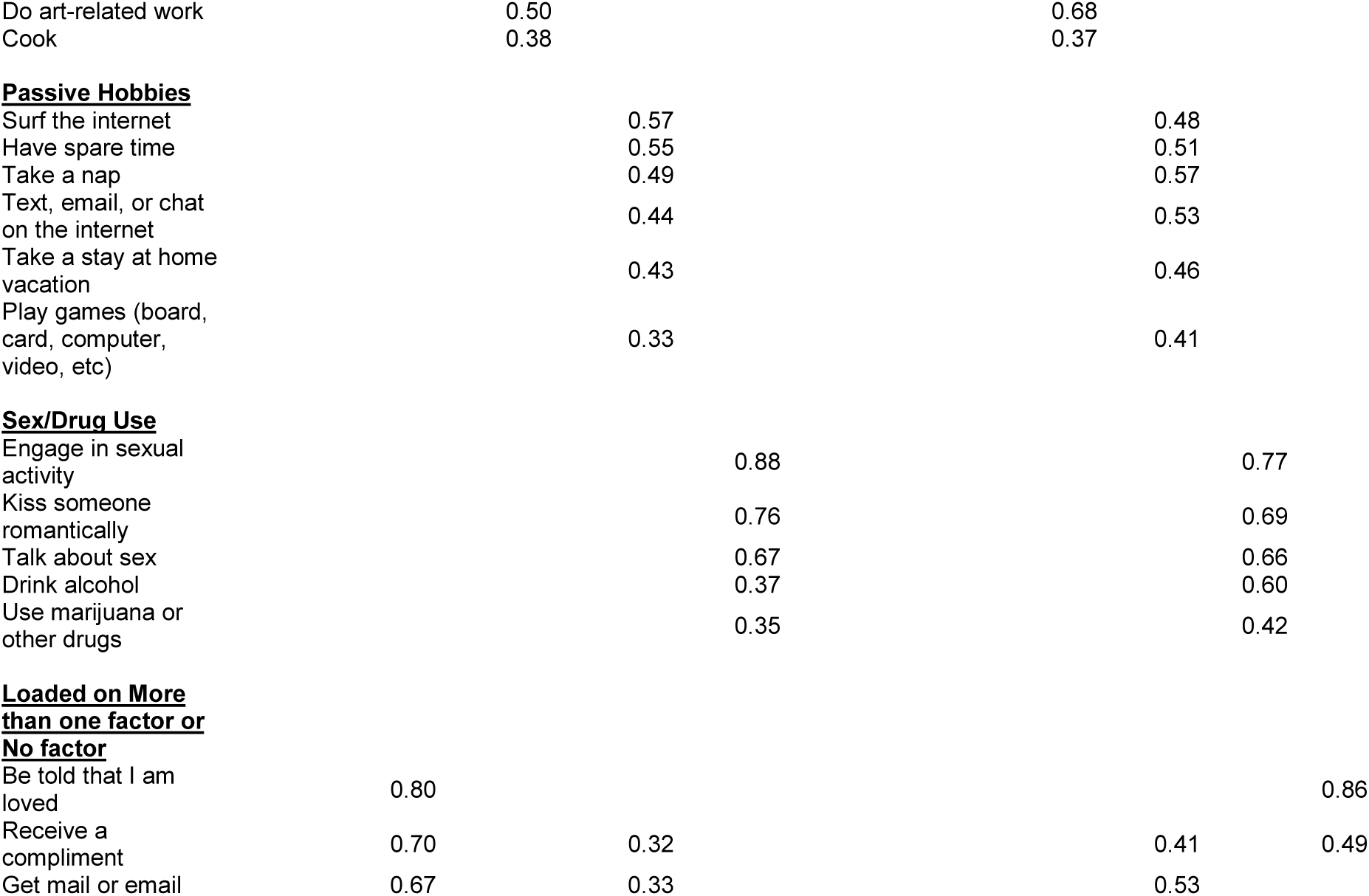
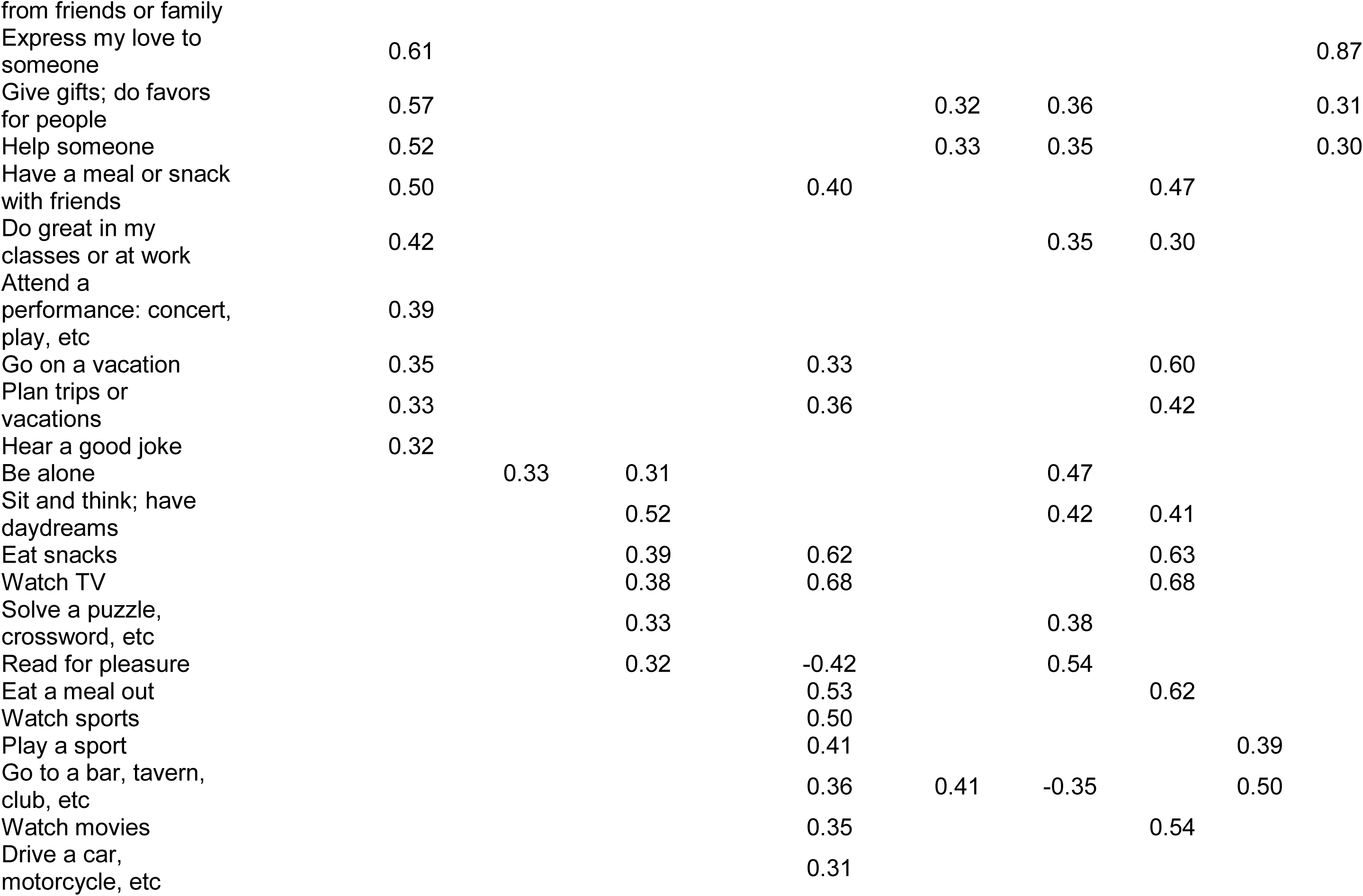
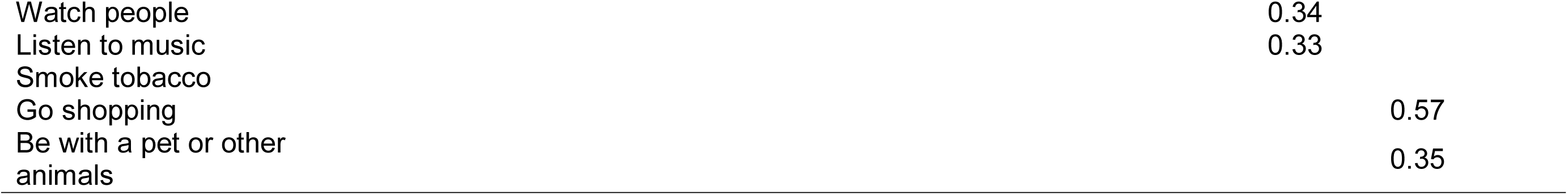
Factor Analysis

**Appendix Table 2.** Overall and Factor Scores, by Sex, Ethnicity/Race, Education, and Age

|  | *n* | Overall REI<br>Enjoy | Socializing | Active<br>Hobbies | Passive<br>Hobbies | Sex/Drug<br>Use |
| --- | --- | --- | --- | --- | --- | --- |
|  |  | *M (SD)* | *M (SD)* | *M (SD)* | *M (SD)* | *M (SD)* |
| Sex |  |  |  |  |  |  |
| Male | 218 | 3.6* | 3.6** | 3.5 | 3.4)*** | 3.3*** |
| Female | 222 | 3.7 | 3.8 | 3.7 | 3.7 | 3.0 |
| Ethnicity/Race |  |  |  |  |  |  |
| Non-Latino White | 385 | 3.6 | 3.7 | 3.6 | 3.5 | 3.1* |
| Other | 55 | 3.7 | 3.9 | 3.7 | 3.6 | 3.3 |
| Education |  |  |  |  |  |  |
| High school or less | 104 | 3.7 | 3.8 | 3.7 | 3.6 | 3.1 |
| Some college or more | 335 | 3.6 | 3.7 | 3.6 | 3.5 | 3.1 |
| Age |  |  |  |  |  |  |
| 18-31 | 108 | 3.7*** | 3.8* | 3.6 | 3.6 | 3.4*** |
| 32-43 | 109 | 3.7 | 3.74 | 3.7 | 3.6 | 3.3 |
| 44-54 | 113 | 3.6 | 3.65 | 3.6 | 3.5 | 3.03 |
| ≥55 | 110 | 3.5 | 3.52 | 3.5 | 3.4 | 2.79 |

\* $p \leq 0.05$
\*\* $p \leq 0.01$
\*\*\* $p \leq 0.001$

